# Local Subspace Pruning (LSP) for Multichannel Data Denoising

**DOI:** 10.1101/2022.02.27.482148

**Authors:** Alain de Cheveigné

## Abstract

This paper proposes a simple algorithm to remove noise and artifact from multichannel data. Data are processed trial by trial: for each trial the covariance matrix of the trial is diagonalized together with that of the full data to reveal the subspace that is – locally – most eccentric relative to other trials. That subspace is then projected out from the data of that trial. This algorithm addresses a fundamental limitation of standard linear analysis methods (e.g. ICA) that assume that brain and artifact are linearly separable within the data. That assumption fails if there are more sources, including noise and brain sources, than data channels. The algorithm captitalizes on the fact that, if enough of those sources are temporally sparse, linear separation may succeed locally in time. The paper explains the rationale, describes the algorithm, and evaluates the outcome using synthetic and real brain data.

## 1 Introduction

Electrophysiological and imaging data are typically contaminated by noise and artifacts that corrupt or mask more subtle signals from the brain. Artifacts and noise may come from environmental sources, potentials that arise at the interface between electrode, electrolyte, and skin, and physiological phenomena such as muscle or ocular activity. Irrelevant brain sources too may compete with brain activity of interest.

Fortunately, multichannel data (from multiple electrodes, sensors, pixels or voxels) can be leveraged to form linear combinations (spatial filters) that attenuate the noise and enhance the useful signal. Many techniques are available to find good sets of coefficients for these filters, some based on sensor geometry (e.g. gradient or Laplacian), others using data-driven algorithms such as Independent Component Analysis, ICA (Hyvärinen et al., 2009)), Common Spatial Patterns, CSP, (Koles et al., 1990)), or others (Parra et al., 2005) to find the necessary weights. Unfortunately, whatever the algorithm, the effectiveness of linear filtering is limited if sources are not *linearly separable*. In particular, if there are more sources than data channels, there may exist *no* set of filter coefficients that that allows all interfering sources to be suppressed.

Nonetheless, if some of the sources are temporally sparse, the number of sources active on each trial might be smaller. For example, movement artifacts might occur on just a few trials, or muscle artifacts might likewise be sporadic. In that case, trial-specific analysis might be more effective than an analysis common to all the data. The present method capitalizes on this idea.

While the idea of trial-specific analysis is attractive, it does raise several concerns: conceptual complexity, algorithmic complexity, risk of overfitting because each trial contains a small amount of data, and risk of bias if the analysis on trials of one condition differs from that for trials of another condition. These concerns are partly mitigated by the focus on *denoising*, as long as noise is not systematically biased to one condition relative to others.

The method is designed as a preprocessing step prior to data analysis. This makes it widely applicable, and relatively simple to understand. At each step the algorithm removes from each trial the particular component that is most *dissimilar* from all other trials. Less eccentric dimensions, which span the bulk of the data, are untouched. Activity that recurs on multiple trials is less likely to be unaffected.

Other methods have been proposed that also leverage the non-stationary correlation structure of the data. These include *adaptive beamforming* (Sekihara et al., 2006; Quraan and Cheyne, 2010; Wong et al., 2018), *adaptive CSP* (Samek et al., 2012; Vidaurre et al., 2015; Jiang et al., 2020; Yu et al., 2020), *sparse time artifact rejection (STAR)* (de Cheveigné, 2016), *inpainting* (de Cheveigné and Arzounian, 2018), and others. Relations between approaches are reviewed in the Discussion.

## Methods

This section specifies the assumptions and describes the algorithm.

### 1.1 Assumptions

#### Data Model

The raw data consist of a matrix **X** of dimensions *T* (time) × *J* (channels) that groups the time series recorded from *J* sensors or electrodes. This matrix is assumed to reflect the activity of *I* sources (both brain and artifact) via a mixing matrix **M** of size *I* × *J*:

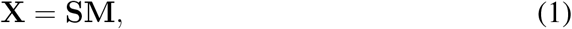

where **S** represents the time series of source activations. **X** is observed, **S** and **M** are unknown. Blind source separation methods (e.g. ICA) produce an unmixing matrix **U** that yields a matrix **Y** of “component” time series:

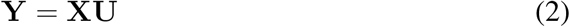

Ideally, one would like the product **MU** to be the identity matrix, so that all sources are recovered, but that goal is unrealistic if the number of sources exceeds the number of observations (*I* > *J*). A more realistic goal is to find a spatial filter **u** that cancels the major sources of interference to obtain a signal **y** reflecting activity of of interest:

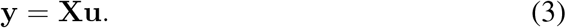

Given *J* channels, up to *J* − 1 sources can be canceled in this way *but no more*. This is a fundamental limit: a spatial filter cannot reject more interfering sources than the number *J* of channels, minus one. This is true whatever the method used to find the filter (ICA, CSP, decoding, etc.).

#### Temporal sparsity

Fortunately, many sources are temporally. Based on this assumption, we can hope to cancel more interfering sources than channels by applying different spatial filters to different time intervals. For definiteness and simplicity of exposition (and because the situation arises commonly), data are assumed to have been re-organized by trials in the form of an array **X**_*n*_ of matrices of size *T*_*n*_ (time) × *J* (channels) × *N* (trials). Temporal sparsity of a source, in this case, means that it is active during one or a few trials and quiescent in all others.

If trial *n* contains at most *J* − 1 interfering sources, those sources can be removed by applying the appropriate trial-specific filter **u**_*n*_. For *N* trials to a total of *N* (*J* − 1) interfering sources might be handled in that way. That ideal unlikely to be attained (or needed) in practice: a realistic and worthy goal is to zap just a few more than the *J* − 1 interfering sources allowed by standard methods, and thus improve the quality of analysis or decoding.

### 1.2 Algorithm

The algorithm operates by repeatedly applying the Joint Diagonalization (JD) algorithm (Fukunaga, 1972; de Cheveigné and Parra, 2014), also known as as DSS (Denoising Source Separation Särelä and Valpola, 2005) or CSP (Common Spatial Patterns Koles et al., 1990), to maximize the ratio of power of each trial relative to the rest of the data. This is obtained by jointly diagonalizing the covariance matrix **C** of the entire data, and the covariance matrix **C**_*n*_ of the trial. Specifically:

1. For trial *n* = 1... *N*:

a. Calculate the covariance matrix **C**_*n*_ of that trial.
b. Calculate the covariance matrix **C** of the entire data.
c. Solve the generalized eigenvalue problem for **C**_*n*_ and **C**, record the largest eigenvalue.
2. If the largest eigenvalue over trials is less than a threshold Θ, terminate, else:

a. Select the corresponding trial *n* and calculate the transform matrix **U**_*n*_ that jointly diagonalizes **C**_*n*_ and **C**.
b. Calculate its pseudo inverse 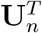, delete the first column of **U**_*n*_ and the first row of 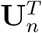, and form their product to obtain the denoising matrix **D**_*n*_.
c. Apply **D**_*n*_ to **X**_*n*_ to denoise trial *n*.

This sequence is repeated *M* times. Parameters are Θ and *M*.

Joint diagonalization (step 2 a) produces a matrix **U**_*n*_ that transforms the data such that the first component has the highest possible ratio of power within trial *n* relative to other trials. Deleting the first column of **U**_*n*_ effectively cancels that component, and back projection results in data that are identical to the original apart from the deleted component. The denoising matrix **D**_*n*_ subsumes projection, deletion, and back-projection. The other *J* − 1 components are not affected, nor are the other trials, although they might be affected on a different iteration.

Many experiments use trials to repeatedly sample brain activity of interest. Such activity, that recurs on most trials, is unlikely to be removed. This assumption is reasonable for typical stimulus-evoked or stimulus-induced activity from spatially stable sources, but it might not hold for activity that is sporadic or spatially diverse (e.g. epileptic activity). The algorithm might not be appropriate for such data.

The computation cost is dominated by eigenvalue decomposition 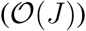, which is repeated *M N* times unless the algorithm terminates early. Optionally, the cost can be reduced by by applying the same matrix **D**_*n*_ to a group of trials for which the trial-to-total power ratio of the first column of **XU**_*n*_ is above some threshold Θ_2_ < Θ. The algorithm then has a third parameter Θ_2_.

For the algorithm to be effective, five conditions must be met: (a) artifacts must have a spatial signature distinct from sources of interest, (b) they must be temporally sparse, (c) their number must not exceed *J* on any trial, (d) sources of interest must be recurrent, and (e) their spatial signature must be stable. If those conditions are not all met, the algorithm may not be effective.

### 1.3 Evaluation datasets

The algorithm is evaluated using synthetic data, real electroencephalogram (EEG) data, and hybrid data (synthetic target embedded in real EEG noise). The advantage of using a synthetic target is that the performance of the algorithm can be accurately quantified. The busy reader might want to skip this section and come back for details as needed.

#### Synthetic dataset A

A data matrix **X** of size *T* = 1000 samples × *J* = 30 channels × *N* = 50 trials is obtained as the sum of one target (a pulse-shaped sinusoid) mixed with 50 transient Gaussian noise sources, each active during one trial and quiescent during all others. The target is mixed into the data via a 1 × 30 mixing matrix with random coefficients, and the interference via 50 × 30 mixing matrix with random coefficients, such that the signal-to-noise ratio (in power) is SNR=0.001. Since there are more sources than channels (one target, 50 interference) the sources are *not* linearly separable within the data.

#### Synthetic dataset B

A data matrix **X** of size *T* = 1000 samples × *J* = 30 channels × *N* = 50 trials is obtained by mixing one target source (a pulse-shaped sinusoid) with 20 stationary Gaussian noise sources. The target is mixed into the data via a 1 × 30 mixing matrix with random coefficients, and the interference via 50 × 30 mixing matrix with random coefficients. Since there are more channels than sources, the sources *are* linearly separable.

#### Synthetic dataset C

Dataset **C** is the same as dataset **B**, with the addition of 50 transient sources as in Dataset **A**, resulting in *I* = 71 sources (one target, 20 stationary noise, 50 transient noise). Since there are now more sources than channels, the sources are again *not* separable within these data.

#### Hybrid dataset D

This data set uses a synthetic target mixed into a background of real EEG. A data matrix **X** of size *T* = 2100 samples × *J* = 30 channels × *N* = 50 trials was obtained by adding the target (a pulse-shaped sinusoid) to one channel of a 30-channel sample of real EEG. The EEG data were taken from a pilot recording for an unrelated study, and cut into “trials”. The EEG background is is assumed to include both stationary and transient sources, thus challenging both LSP and standard linear analysis. Since the source is known, however, performance can be quantified accurately.

#### EEG dataset E

Data were taken from a publicly available dataset (https://zenodo.org/record/3618205) described in a published study (Fuglsang et al., 2020). In brief, 44 subjects listened to 180 repetitions of a 1 kHz tone of duration 100 ms (the study involved other stimuli and tasks not considered here). EEG were recorded at 512 Hz sampling rate on 66 channels (64 scalp and 2 periocular) using a BioSemi Active II system. After mean removal, detrending, and 50 Hz artifact removal (de Cheveigné, 2019), peri-stimulus epochs were excised and organized as a matrix of size *T* = 132 samples × *J* = 66 channels × *N* = 180 trials. Each channel of each trial was again detrended and its mean removed.

#### EEG dataset F

Data were taken from a pilot recording for a study involving both auditory and visual stimulation, the same as for Dataset **D**. Subjects listened to a train of concurrent audio and visual stimuli, consisting of 50 ms tones (audio) or dots (visual). At randomly-spaced instants, the train transitioned from a disorderly state (frequencies and positions chosen at random) to an orderly state (a proportion of tone frequencies and/or dot ordinates increasing regularly). Subjects were asked to detect those events. Brain responses were recorded at 512 Hz sampling rate from 64 scalp electrodes, and the data were organized by trials (time-locked to the stimulus transition) as a 3D matrix of size *T* = 871 samples × *J* = 64 channels × *N* = 672 trials. The JD algorithm was applied to suppress eye blink correlates, using the instantaneous power at electrodes closest to the eyes as a mask, as described in de Cheveigné and Parra (2014), and the mean and linear trend of each trial and channel were removed using a robust detrending algorithm.

### 1.4 Evaluation methods and metrics

LSP is intended as a preprocessor to improve the outcome of other analysis methods, and its effectiveness may be judged as an increment in success rate of those methods. This section briefly describes those analysis methods. The busy reader might want to skip this section and come back for details as needed.

#### Trial Average

If the target signal repeats on each trial, but background activity does not, averaging improves the SNR by a factor of *N*, the number of repeats. Using the trial mean 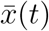 as an approximation of the “true” target, the noise can be approximated as 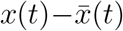 where 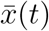 is the trial average replicated over all trials. The SNR of the raw data can then be estimated as 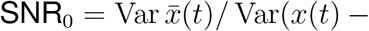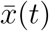. The SNR of the average over trials is then estimated as SNR= *N* SNR_0_.

#### Linear Component Analysis

The multichannel nature of EEG or MEG can be leveraged to form linear combinations of channels with better SNR. The appropriate coefficients can be found using data-driven component analysis algorithms. Here, the JD algorithm is used with a bias designed to maximize repeatable power, i.e. the ratio of power of the trial mean over that of the raw data. This algorithm (denoted as JD_*r*_) produces *J* components, the first of which has the highest SNR as defined above. That component can then be averaged over trials, yielding a better SNR than averaging any of the raw channels (de Cheveigné and Simon, 2008; de Cheveigné and Parra, 2014).

The JD_*r*_ solution is obtained by jointly diagonalizing the covariance matrices of the raw data and of the trial-averaged data. It is also possible to jointly diagonalize the covariance matrices of two different time intervals or conditions (denoted as JD_*t*_), in which case the method is identical to the well-known Common Spatial Patterns (CSP) method (Koles et al., 1990). Finally, it is also possible to jointly diagonalize the covariance matrices of raw and filtered data (for example high-pass) to reveal components with particular spectral characteristics (denoted as JD_*s*_). All three variants of the algorithm are employed here.

#### Quadratic Component Analysis (QCA)

QCA addresses the situation where a target is repeatable in *power* (induced response). It aims to find a spatial filter that maximizes the SNR of such a target. For that purpose, all *cross-products* 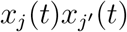 of channels are formed, two-by two, and JD_*r*_ is applied to the resulting matrix of cross-products (de Cheveigné, 2012). The rationale is the following: suppose that *s*(*t*) is a weak source mixed with others and that it can only be recovered as *y*(*t*) = **uX** where **u** is a spatial filter the coefficients of which are unknown. The instantaneous power of the recovered component can be expanded as the quadratic form 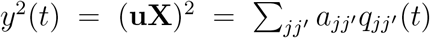, where 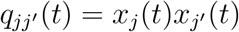 is the instantaneous product between columns *j* and *j*′ of **X**, and 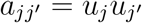. JD*_r_* can then be applied to the matrix of distinct cross-products, of size *T* × *J* (*J* − 1) × *N*, yielding a matrix **Z** of “quadratic components” (QC). The first column *z*_1_(*t*) of **Z** is the most reproducible QC.

While the most reproducible QC might be of interest, one usually wants to find a spatial filter to extract the neural source(s) behind it. Unfortunately, whereas every squared component *y*(*t*)^2^ is the sum of crossproducts of channel waveforms, the converse is not true. However, thanks to the spectral theorem, it is possible to express a quadratic component *z*_*k*_ (*t*) as a weighted sum of squared linear components *y*_*k*1_(*t*)^2^, *y*_*k*2_(*t*)^2^,... etc. Collectively, these linear components can be understood as spanning a subspace of the data that contains reproducible power (de Cheveigné, 2012).

The number of QCs increases rapidly (*J* (*J* − 1)) with the number *J* of data channels. To limit computational cost, and reduce overfitting, dimensionality of the data can be reduced to *J* ′ < *J*, for example by applying PCA and truncating the series of PCs before applying QCA. It is also useful to temporally the cross-product time series 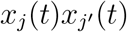 to attenuate irrelevant fine structure, and downsample to reduce memory cost.

#### Metrics

For synthetic data, where the source *s*(*t*) is known, performance can be quantified by the metric 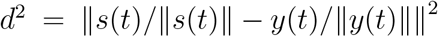, where *y*(*t*) is the outcome of analysis. Its value ranges from 0 (perfect retrieval) to 2 (source and estimate uncorrelated). For real data, the source is unknown, but it can be approximated based the outcome of data analysis (e.g. the trial average of the best JD_*r*_ component for an evoked response).

## 2 Results

### 2.1 Synthetic and hybrid data

#### LSP effectively removes transient sources

Dataset **A** consists of a repetitive target (Fig. 1 a) on the background of transient noise sources, each active during one trial and quiescent during others (Fig. 1 b). The mix is heavily dominated by noise (Fig. 1 c) that the LSP algorithm largely removes (Fig. 1 d). A standard linear analysis methods would have failed because the sources are not linearly separable.

**Figure 1:**
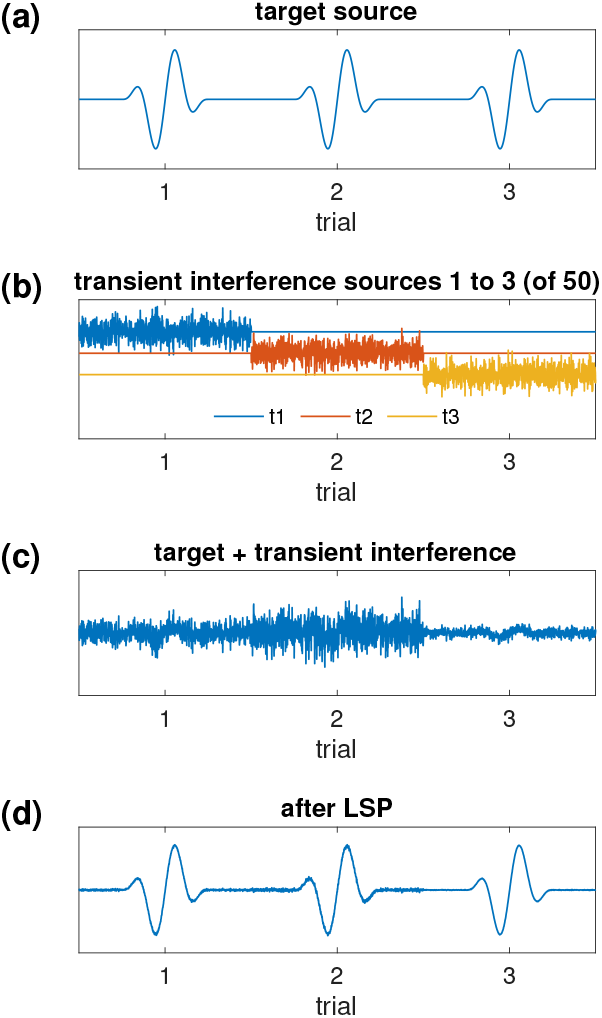
LSP applied to synthetic data. The target source (a) is mixed with 50 interference sources (b, 3 sources shown) yielding a mixture dominated by interference (c, one channel shown) with SNR = 10−3 in power. Applying LSP recovers the target. Each plot shows a concatenation of the first 3 trials (of 50).

#### Linear analysis applied to linearly-separable data

Dataset **B** consists of a repetitive target (Fig. 2 a) on the background of 20 stationary Gaussian noise sources (three of which are shown in Fig. 2 b). The 30-channel mix is noisy (Fig. 2 c), but the SNR can be improved by applying JD_*r*_ then averaging over trials (Fig. 2 d). Linear analysis (JD_*r*_) is successful because the sources are linearly separable. This example is not intended to illustrate LSP, but rather to provide backdrop for the next simulation.

**Figure 2:**
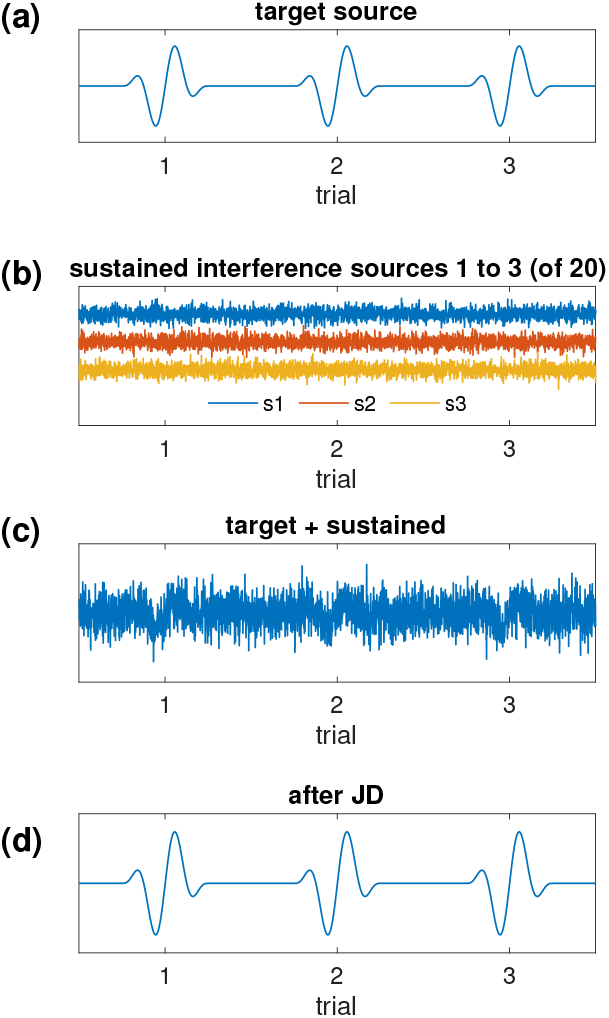
Linear analysis of linearly separable synthetic data. The target source is mixed with 20 stationary interference sources (b, 3 sources shown) yielding a mixture dominated by interference (c, one channel shown). Applying JD_r_ recovers the target (d). The overall SNR is 10−3 in power.

#### LSP helps isolate a synthetic target from a synthetic background

Dataset **C** consists of a repetitive target (Fig. 2 a) on the background of 20 stationary Gaussian interference sources (Fig. 2 b) and 50 transient interference sources (Fig. 1 b). The 30-channel mix is noisy (Fig. 3 a). Applying JD_*r*_ alone is ineffective because the data are not separable (Fig. 3 c, compare with Fig.2 d). LSP alone is also ineffective because it only removes the transient sources, not the stationary sources (Fig. 3 b). However applying first LSP then JD_*r*_ is effective (Fig. 3 d). LSP boosts the effectiveness of linear analysis method by pruning transient sources, so that the remaining sources are separable.

**Figure 3:**
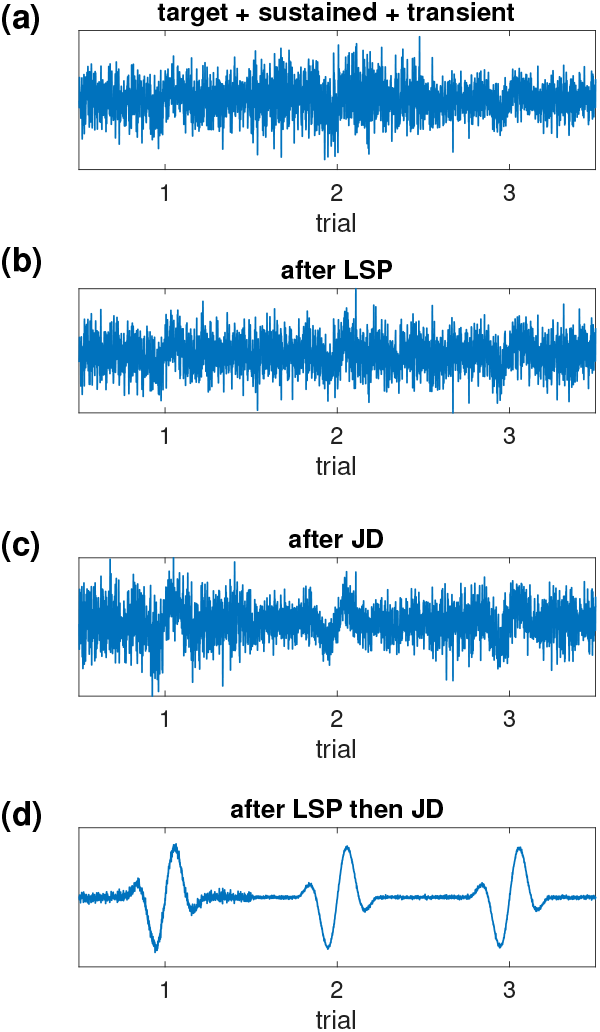
Combination of LSP and JD_r_ applied to synthetic data. The target source (Fig. 2a) is mixed with 20 stationary interference sources (Fig. 2a) and 50 transient sources (Fig. 2b), yielding a non-separable mixture dominated by interference (a). Applying LSP alone improves the SNR slightly (b), as does JD_r_ alone (c). Applying both yields a larger benefit (d). Sustained and transient interference sources have the same power, and the overall SNR is 5 × 10^−3^.

The average over trials of the same data is plotted in Fig. 4 (a). The blue line indicates the trial average, and the gray band shows ±2 standard deviations of a bootstrap resampling of this mean. Preceding the trial average by JD_*r*_ only (Fig. 4 b) reduces the variability slightly, as does preceding it by LSP only (Fig. 4 c), but preceding it by both offers a greater improvement (Fig. 4 d). JD_*r*_ by itself is ineffective because the data are not linearly separable, LSP by itself is ineffective because it cannot deal with stationary noise sources.

**Figure 4:**
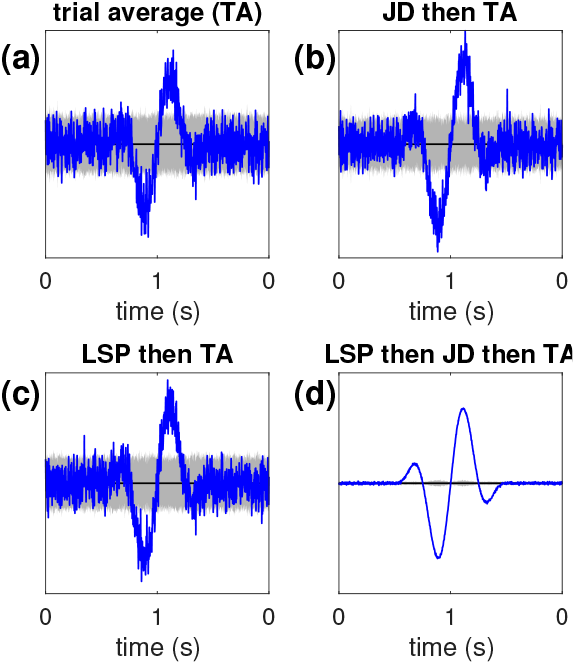
Same as Fig. 3, averaged over trials. The width of the gray band, which represents ±2 SD of the mean over trials, quantifies reliability (narrower is better).

#### LSP helps suppress a real EEG background

Dataset **D** consists of a repetitive synthetic target (Fig. 2 a) on the background of real EEG (Methods). EEG differs from Gaussian noise by its complex serial and cross-channel correlation structure. At this SNR (0.025) the mixture is strongly contaminated by noise (Fig. 5 a, b). JD_*r*_ (c) by itself or LSP by itself (d) are both relatively ineffective, but applying them in succession is more successful (e).

**Figure 5:**
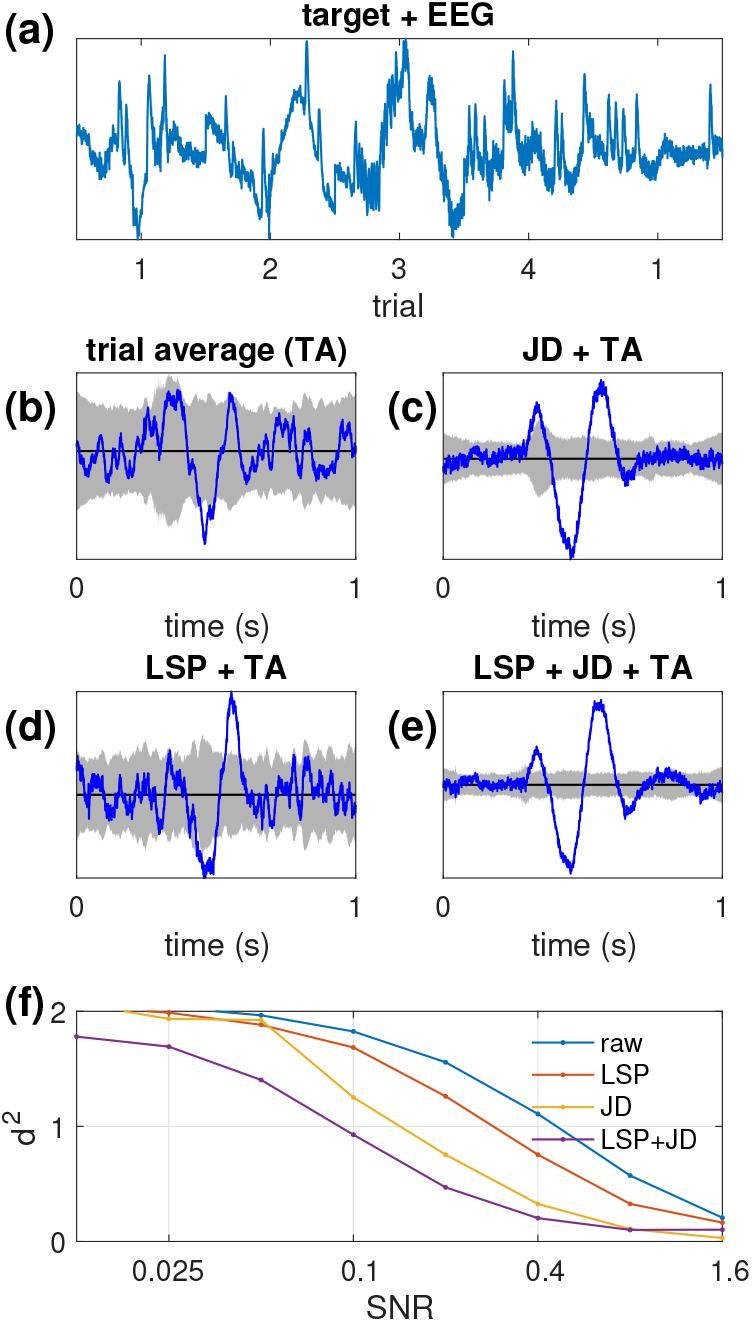
Combination of LSP and JD_r_ applied to hybrid data. A repetitive synthetic target (Fig. 2a) is mixed with EEG at an unfavorable SNR (0.025). The noisiness of the trial average is reflected by the width of the gray band in b. Applying JD_r_ reduces variability (c), as does applying LSP (d), but the greatest improvement is obtained by applying both (e). This benefit holds over a wide range of SNR (f)

Since the target is known, it is easy to quantify the mismatch between the original and recovered target. Figure 5 (f, left)) shows the error (*d*^2^) as a function of SNR for the unprocessed and processed data. LSP by itself offers little benefit, but JD_*r*_ by itself is beneficial across most of the range of SNR. Preceding it with LSP offers an additional increment of performance, particularly at small SNR.

The benefit is less marked than with a synthetic background, presumably because a proportion of noise sources are not “sparse” in the sense required by the algorithm (active on only one or a few trials). Nonetheless, a benefit is found over a wide range of SNRs, particularly at unfavorable SNR.

### 2.2 Real EEG data

The question that remains is whether LSP can demonstrate benefit to extract activity of interest from real brain data. Anticipating, the answer is nuanced.

#### LSP offers little benefit in an evoked-response paradigm

Dataset **E** consists of EEG recorded in response to 180 repetitions of a 1 kHz tone of duration 100 ms from 4 subjects. The data, recorded from 64 scalp and 4 peri-ocular electrodes, were preprocessed to remove 50 Hz noise, slow drift, and ocular artifact, and organized by trials in a 3D matrix (Methods). The data were variously processed with JD_*r*_ to enhance repeatable activity, by LSP, and by a cascade of LSP and JD_*r*_. Figure 6 shows that LSP provides *no* benefit in this case.

**Figure 6:**
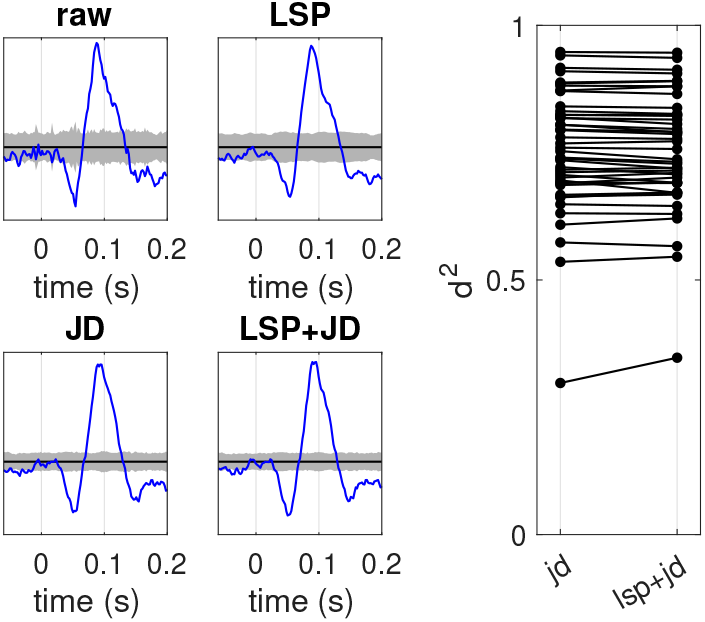
Real EEG, evoked auditory response. LSP offers no benefit for this task.

This lack of benefit is sobering. It could mean either that temporally-sparse artifacts are not prominent in these data (they do not impact the dimensions that carry the auditory response or the ability of JD_*r*_ to enhance these dimensions) *or* that the auditory response itself is somewhat sparse and is thus attenuated by LSP.

#### LSP helps reveal a gamma-band auditory evoked response

This example also uses dataset **E**. The data of one subject (subject 2) were differentiated twice and submitted to a LSP (threshold = 5, iterations = 30) followed by JD_*r*_ to find the maximally repeatable linear combination of channels.

Figure 7 (a) shows a raster plot of the time course of this component over the first 100 trials. On almost every trial, the stimulus onset triggers a burst of pulse-like events with a latency of approximately 18 ms that fades after about 20-30 ms until a less reliable cluster roughly contemporary with the offset of the 100 ms tone.

**Figure 7:**
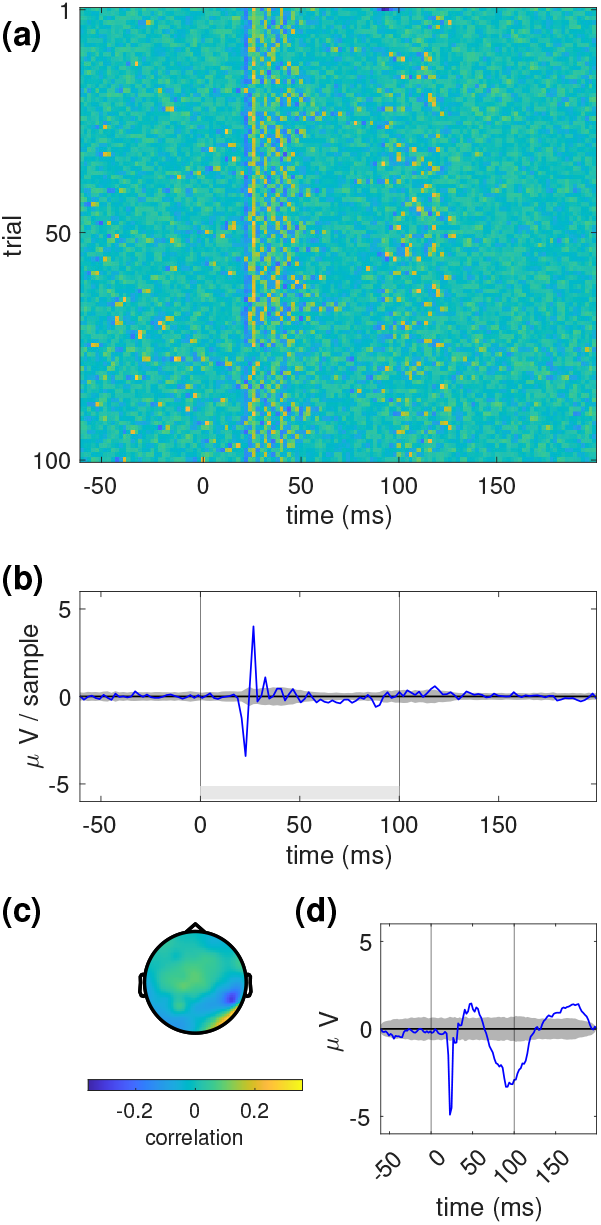
Real EEG, evoked auditory gamma response. (a) Raster plot of second derivative of first JD_r_ component for each trial. (b) Average over trials (blue) and two standard deviations of a bootstrap resampling of the mean. (c) Topography of the corresponding component. (d) Average of the same component twice integrated.

The initial pulse of each burst appears to be tightly locked to the stimulus onset, with a jitter on the order of one sample (∼2 ms), reflected by a prominent pulse in the trial average (Fig. 7b, blue). Subsequent pulses are less well aligned, as reflected by the wide gray band in the trial average. Visually, the pulses seem regularly spaced, but spectral analysis yields a rather wide peak at around 150 Hz (not shown), suggesting that the inter-pulse interval is rather variable. Interestingly, the second, fainter burst seems to start *before* the tone offset, at least on some trials. A few isolated pulses occur before the stimulus, but rarely after it. Integrating this response twice (to reverse the effect of differentiation) reveals that these sharp pulses ride on a slower oscillatory response (Fig. 7 d).

This component is maximally correlated with a pair of right parietal electrodes (P8, P10, Fig. 7c). It is tempting to attribute it to a gamma band response from primary auditory cortex, given the short latency, which would be remarkable because such responses have mainly been observed from invasive recordings (Edwards et al., 2005), rather than EEG. However, an alternative account is that it reflects myogenic activity from the post-auricular reflex (PAR), the fastest exteroceptive reflex with a disynaptic or trisynaptic pathway (Hackley, 2015).

Regardless the interpretation, the conclusion is that LSP enables this weak response to be isolated. The response is observed with high temporal resolution on a trial-by-trial basis (Fig. 7 a), in contrast to most reports of gamma-band activity that rely on spectrographic representations (e.g. Sinai et al., 2009). Spectral analysis entails loss of temporal resolution (de Cheveigné and Nelken, 2019), so spectrogram representations are temporally imprecise.

#### LSP helps reveal a stimulus-induced response

The previous examples targeted a stimulus-evoked response, time-locked to each trial. In contrast, this example targets a stimulus-*induced* response for which the envelope (or instantaneous power) is time-locked, but not the waveform. Induced responses are not amenable to trial-averaging, and cannot be enhanced by JD_*r*_.

Instead, a different technique is used. In QCA, all cross-products between channels are formed, and JD_*r*_ is then applied to find the most repeatable quadratic form (linear combination of cross-products), from which a spatial filter can be derived to enhance the source of induced activity. Since quadratic quantities (squares and cross-products) are very sensitive to high-amplitude outlier values, this analysis is rather brittle and hard to apply to real data. LSP can help by removing trial-specific artifacts.

This example uses Dataset **F**, that consists of 64-channel EEG responses recorded in response to concurrent trains of auditory and visual stimuli (Methods). Subjects detected a change in stimulus characteristics and responded by a button press. After detrending, high-pass filtering, rereferencing to the mean, the data were reduced to 30 dimensions using the SCA algorithm (de Cheveigné, 2021). The LSP algorithm was then applied with parameters *M* =10 (number of passes) and Θ=10 (threshold) to remove transient noise sources.

The QCA algorithm was then applied to the data. All distinct cross-products of channels were formed into a matrix of size *T* =871 × *J* (*J* − 1)= 870 × *N* =672 which was then smoothed by a square window of size 26 samples (approximate period of the 20 ms stimulus) and downsampled by a factor of 26. The mean was removed from each trial, and the JD_*r*_ algorithm was applied to this matrix of cross-products to maximize repeatability. The two most repeatable components (QCs) are shown in Fig.8 (a, left), and the main linear components underlying these QCs are in Fig.8 (a, right), plotted in terms of power relative to stimulus onset. These include a number of components for which power decreases (dotted lines), a response classically known as “event related desynchronization” (ERD), in addition to two components for which power instead *increases* (full lines), a response known as “ event-related synchronization” (ERS),

The 2-dimensional subspace containing these components can be rotated using the spectral version of JD (JD_*s*_, Methods) to maximize their spectral dissimilarity. The resulting two components, labeled HF and LF, have topographies shown in (b) and time-courses shown in (c) for three selected trials. Interestingly, the HF component (blue) consists of spikes that are “sandwiched” between two events of opposite polarity in the LF component (red).

It is tempting to interpret these responses as a stimulus-induced. However, a standard evoked-response analysis time-locked to the button press suggests that both components are instead response-related (not shown). The LF component appears to be a subtle artifact of the BioSemi response box that contaminates the EEG channels, which explains the opposite polarities of the first and second pulse (corresponding button press and button release). The HF component is likely muscular, possibly mediated by a collateral of the motor efferent that effectuates the button press, which would explain why it is sandwiched between LF pulses. Thus, this “stimulus-induced” response is probably merely a combination of response artifacts.

Regardless the interpretation, the point is again that LSP allowed this weak response to be extracted and plotted with high temporal resolution on a trial-by-trial basis (Fig. 8 c).

**Figure 8:**
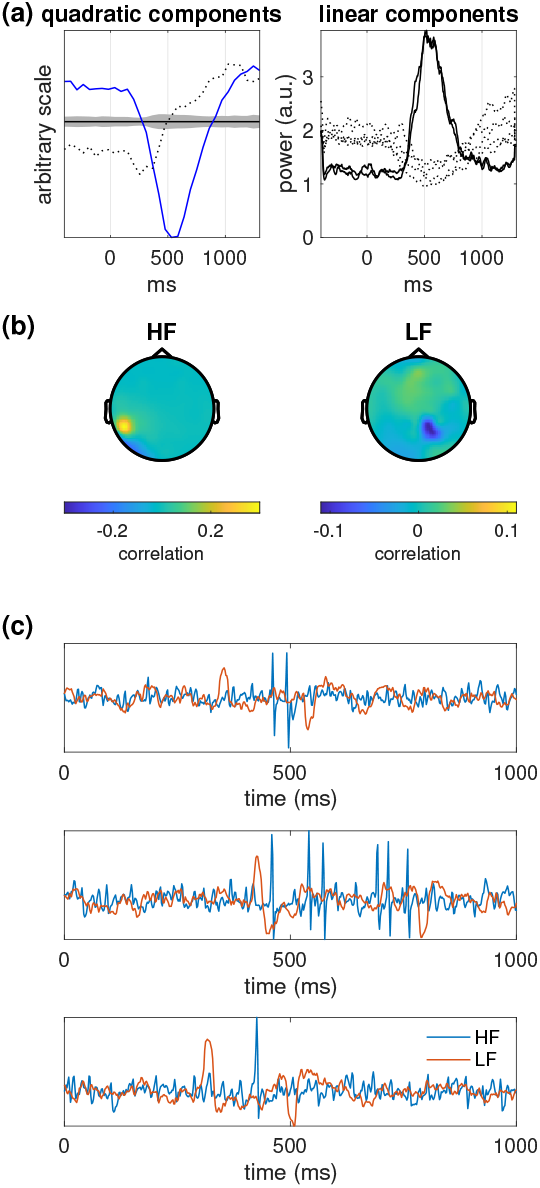
Real EEG, induced response. (a) Left: trial-average (blue) and 2SD of the bootstrap resampling (gray) of the first QC. The trial-average of the second QC is shown as a dotted line. Right: time course of selected linear components corresponding to the selected QCs. The two full lines represent components for which power increased after stimulus onset (ERS), the dotted lines components for which it decreased (ERD). (b) Topographies of the two ERS components after rotation to maximize spectral dissimilarity. HF: high frequency, LF low frequency. (c) Time course of the LF and HF components for three selected trials.

## 3 Discussion

LSP addresses the situation where there are more sources of artifact than channels in the data. In that situation, *no* linear analysis method can fully recover the sources. Simulations with synthetic data show that the principle is sound and the method effective. Evaluation with real EEG data give more nuanced results: in some situations LSP provides no obvious benefit, in others it is effective in enabling linear analysis (JD) to extract weak sources.

LSP is thus mainly useful as a specialized tool, rather than a module in a standardized pipeline. It may be precious in specific situations to discover weak sources that other methods cannot reveal. For example, the LF source in the last example represented about 1% of the power of the best electrode (Fig. 8 b, right), so it would be impossible to discover it without these analysis methods, and certainly not plot it on a trial-by-trial basis (Fig. 8 c).

### Vector space interpretation

A data matrix **X** of size *T* (time) × *J* (channels) × *N* (trials) consists of *TJN* values, i.e. it can be represented as a point within a space of dimension *TJN*. However, linear analysis methods (ICA, JD, etc.) are constrained to form linear combinations of only *J* channels, i.e. they operate within the smaller *J*-dimensional subspace spanned by the channel signals. Sources (brain or artifact) are known only from their projection on this subspace.

If there are more sources than channels (*I* > *J*), the mixing matrix **M** is not invertible and one cannot find an unmixing matrix **U** to recover the sources. The best one can hope for is to cancel the strongest interfering sources. At most *J* − 1 sources can be cancelled in this way. If sources are more numerous, which is likely given the complexity of brain activity and the many sources of artifact (e.g. myogenic activity or contact noise at each electrode), they cannot all be suppressed.

However, if a different matrix **U**_*n*_ is applied on each trial, a larger number of interfering sources might be suppressed, as many as *N* (*J* − 1) in principle. LSP capitalizes on this flexibility.

### Other approaches

A common procedure is to discard artifact-contaminated trials. This might be effective, but it is unnecessary if those trials could be cleaned rather than deleted, and impractical if too many trials are involved.

Several methods address the case where temporally-sparse artifacts affect single channels, or groups of channels. For example the STAR algorithm (de Cheveigné, 2016) reconstructs the waveform of one channel based on the waveforms of the other channels, via a linear model learned from the data. It requires the data to be artifact-free for enough of its duration so that its correlation structure can be estimated, a requirement that is softened in a more recent version (de Cheveigné and Arzounian, 2018). Those algorithms reconstruct missing values based on the correlation structure estimated from the data, but it is also possible to use the sensor geometry to interpolate from neighbours. In every case, the assumption is that the artifact affects a single channel (or small number of channels). In contrast, LSP allows the temporally-local artifact to impact a subspace spanned by multiple channels.

Another approach is to design the data analysis stage to be adaptive. Examples are the adaptive CSP approach of (Jiang et al., 2020), or the stationary CSP approach of (Samek et al., 2012). The latter resembles LSP in that it seeks to distinguish a subspace stationary across trials from dimensions active on specific trials. It is worth noting a parallel with non-linear dimensionality reduction methods (Burges, 2009) that seek to find, and flatten, a low-dimensional manifold within the data.

## Conclusion

This paper described a simple algorithm for removing non-stationary noise sources in the situation where there are more sources than channels in the data. The algorithm addresses a fundamental limitation of standard linear analysis methods (e.g. ICA) that assume that brain and artifact are linearly separable within the data. It proceeds by projecting out from each trial any component that is eccentric (in terms of correlation structure) with respect to the rest of the data. Tested on synthetic and real data, the algorithm proved effective, although its benefit is task dependent. For some tasks it provides no benefit, for other it reveals weak sources that are otherwise invisible.

## Acknowledgements

Thomas Schaffhauser kindly provided pilot data for the simulations, and Malcolm Slaney and Søren Fuglsang provided useful comments on early drafts of this paper. This work was supported by grants ANR-10-LABX-0087 IEC, ANR-10-IDEX-0001-02 PSL, and ANR-17-EURE-0017.

